# Bivalent bispecific CD28-antibodies reinforce T-cell responsiveness and revert anergy/quiescence in patients treated with bispecific CD3-antibodies

**DOI:** 10.64898/2026.03.25.714198

**Authors:** Latifa Zekri, Natalie Köhler, Ariane Metzger, Nisha Prakash, Jonas S. Heitmann, Monika Engel, Timo Manz, Stefanie Müller, Sebastian Hörner, Karolin Schwartz, Melissa Zwick, Ilona Hagelstein, Nadine Brückner, Martin Pflügler, Josef Leibold, Melanie Boerries, Gundram Jung, Helmut R. Salih

## Abstract

Bispecific T-cell engagers stimulating CD3 (TCEs) rapidly gain momentum in oncological therapy. We report that single-agent TCE-treatment of cancer patients induces T-cell hyporesponsiveness including abolished proliferative and lytic capacity. Single-cell RNA-sequencing identified transcriptional features of anergy/quiescence like upregulation of *CBLB* and *BACH2*, and suppression of AP-1-dependent activation programs due to isolated provision of “T cell signal-1”. To restore T cell functionality, we engineered bispecific costimulators (BiCos) which deliver “signal-2” via highly efficient bivalent CD28-binding yet maintain strictly target-restricted activity after binding antigens expressed not only by tumor cells but also the extracellular matrix. Preclinical analyses documented that BiCos potentiate efficacy of single-agent TCE treatment with activity exceeding that of univalent Knob-into-Hole CD28-costimulators and revert T cell hyporesponsiveness. In humanized mice, BiCos induced elimination of established tumors in combination with very low doses of TCE. Finally, BiCos reversed molecular hallmark features of quiescence such as high expression of *BACH2* and *KLF2* in T cells of TCE-treated patients resulting in complete restoration of cellular function, thereby reinstating durable antitumor immunity.

## Introduction

During the 1980s, central receptors governing the function of T cells - the major effector cells of adaptive immunity - were identified and characterized, starting with the T cell receptor (TCR)/CD3 complex, which mediates the antigen-specific “signal 1” for T cell activation. Not much later, CD28 was identified as prototypical costimulatory receptor delivering an essential “signal 2”. Subsequently, additional costimulatory receptors including 4-1BB (CD137) and OX40 (CD134) were identified (1), and it was confirmed that costimulation not only serves to amplify “signal 1”, but also prevents T cell hyporesponsiveness, which occurs after isolated TCR/CD3 stimulation (2, 3). It was also found that so-called immune checkpoints like CTLA-4 and PD-1 transmit “co-inhibitory” signals that restrain T cell activation (4). These discoveries paved the way for therapeutic modulation of T cell immunity. Beyond bispecific T cell engagers stimulating CD3 (TCEs) and the functionally related CAR T cells, immune checkpoint inhibitors completed the arsenal of T cell recruiting strategies that meanwhile have become a cornerstone of oncological treatment (5, 6). A critical distinction between TCEs and CAR T cells is that the latter require signaling domains not only from CD3, but also from costimulatory molecules (CD28 or 4-1BB) to achieve therapeutic efficacy (7). We introduced costimulatory signals to the field of bispecific antibodies (bsAbs) already in the 1980s by demonstrating that combining bsAbs with TAAxCD3- and TAAxCD28-specificity (TAA, Tumor Associated Antigen), achieved superior T cell reactivity against tumor cells and subsequently conducted a clinical trial evaluating such a combination (8, 9). Subsequently, clinical development of tumor-targeted costimulation was severely impeded by the “TGN1412 (Tegenero) incident” in 2006, when a superagonistic CD28 monospecific antibody (mAb) induced life-threatening cytokine release syndrome (CRS) in healthy volunteers (10).

Our efforts to develop optimized bispecific costimulators (BiCos) were reinforced by recent observations during a clinical study with a PSMA-directed TCE. We here report, to our knowledge for the first time, that prolonged TCE single-agent treatment can induce functional hyporesponsiveness of patient T cells due to suppression of activation programs as well as induction of canonical anergy and quiescence genes. To reinforce T cell function and taking into account that bivalency dramatically enhances the agonistic activity of CD28 antibodies (11), we developed BiCos that, in contrast to so far available constructs, comprise bivalent TAA- and CD28-binding parts while maintaining strictly tumor-restricted activity. As targets we selected endoglin and B7-H3 that are expressed not only on tumor cells but also on the extracellular matrix (ECM) of tumors including the neovasculature. We further reasoned that using two different TAAs with non-overlapping “off-tumor expression” on healthy tissues would provide an additional layer of specificity/safety for rational combinations of TCE and BiCos, as the latter require “signal 1” and thus activation by TCE to mediate their costimulatory function. In this work, we introduce that providing signals via CD28 by using our bivalent BiCos can prevent and also revert T cell hyporesponsiveness occurring upon TCE-single agent-treatment by reinstating proliferative cell-cycle programs as well as enhancing effector signatures and activation-associated pathways while reverting hallmark features of T cell anergy/quiescence, resulting in profoundly increased therapeutic T cell antitumor immunity.

## Results

### Treatment with single agent TCE induces T cell hyporesponsiveness

In an ongoing Phase 1a trial, patients with metastatic castration-resistant prostate cancer (mCRPC) showed a rapid but transient decline of PSA levels upon treatment with our PSMA-directed TCE CC-1 (12) as single agent (NCT04104607). Ex vivo analyses using PBMC of these patients obtained before and shortly after treatment (Figure 1A) revealed a dramatic loss of responsiveness to CD3-stimulation with regard to induction of T cell activation, proliferation, cytokine secretion and cytolytic activity despite comparable CD4⁺ and CD8⁺ T cell counts (Figure 1, B and C; Supplemental Figure 1, A and B). Longitudinal ex vivo analyses revealed that patient T cells partially recovered from this hyporesponsive state after two-week cessation of treatment (Figure 1D, compare e.g. day 7 versus day 22; Supplemental Figure 1C), but overall remained profoundly impaired (with substantial inter-patient variability) as revealed by weakened responses to TCE treatment (Figure 1D, compare day 1 versus days 22, 43 and 64; Supplemental Figure 1C) as well as declining peak serum levels of soluble IL-2 receptor (sIL-2R) upon increasing numbers of treatment cycles (Figure 1E and Supplemental Figure 1D). Hyporesponsive T cells displayed neither a significant upregulation of exhaustion markers (e.g., PD-1: *P* = 0.7 in CD4⁺ T cells and *P* = 0.5 in CD8⁺ T cells) nor a phenotype associated with senescence or terminal differentiation (PD-1, CD57, KLRG1, loss of CD27/CD28) (Supplemental Figure 1, E and F). Single-cell RNA-sequencing (scRNA-seq) of PBMC samples from patients obtained prior to and after TCE treatment identified seven distinct immune cell clusters, including CD4^+^ T cells, CD8^+^ T cells, NK cells, B cells, CD16^+^ monocytes, CD14^+^ monocytes and dendritic cells (Figure 1F). Comparative scRNA-seq of paired day-1 and day-8 samples revealed that TCE-treatment induced a coordinated transcriptional program characteristic for T cell anergy/quiescence: upregulation of regulatory metabolic factors and anergy/quiescence related genes including *CBLB*, *BACH2*, *FOXP1*, *PTPN22*, *LAG3*, *LDHA* and *UCP2*, and downregulation of immediate-early activation genes (*FOS*, *FOSB*, *JUN*, *JUNB*, *DUSP1*, *CD69*) (Figure 1G). This gene expression pattern is consistent with a BACH2-driven quiescent state characterized by repressed AP-1 activation programs and anergy-like inhibitory features, including increased CBLB expression (13, 14). In line, gene set enrichment analysis (GSEA) revealed significant enrichment of anergy-related gene sets (NES = 1.633, *p*<0.0001 and FDR<0.0001 in CD4⁺ T cells and NES = 1.420, *p*<0.0001 and FDR<0.0001 in CD8⁺ T cells) in TCE-exposed T cells (Figure 1H and Supplemental Figure 2). Subclustering and embedding density analysis (Figure 1, I and J) confirmed that CD8⁺ T cells shifted from effector-memory into dysfunctional terminal effector-like states, whereas CD4⁺ T cells transitioned from an early-activated/central memory phenotype into a quiescent/resting state. The observed subcluster distributions are consistent with altered differentiation patterns under chronic, imbalanced stimulation (15).

**Figure 1.**
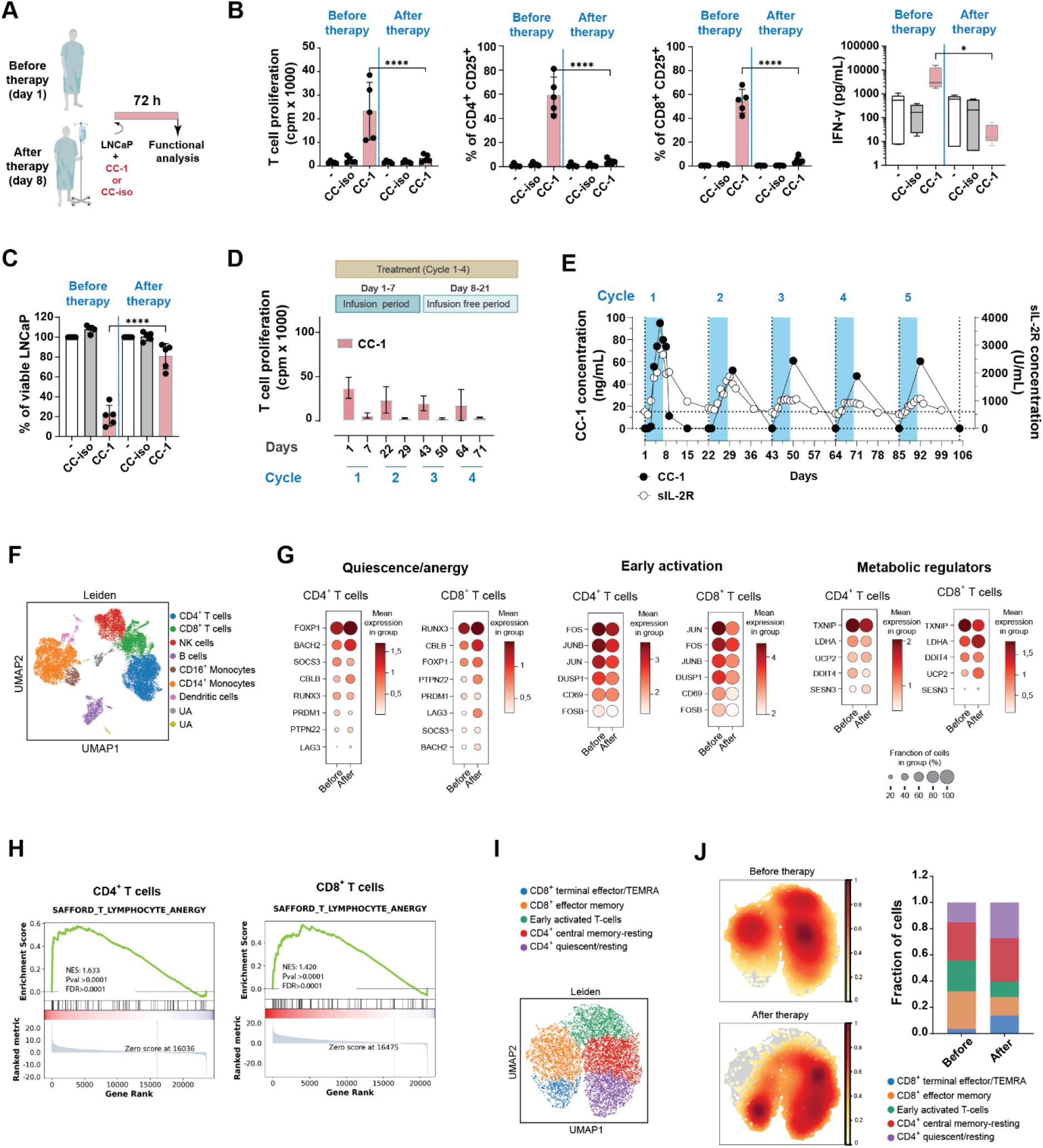
Functional impairment and transcriptional reprogramming of patient T cells upon TCE single agent therapy. (**A**) PBMC (n=5) were collected from patients with mCRPC before and after the first treatment cycle (day 1 and day 8, respectively) and 100,000 cells of either time point were incubated with LNCaP cells (E:T ratio 1:1) for 72 h in the presence of CC-1 or an isotype control antibody (CC-iso) (1 nM each) followed by functional analysis. (**B-C**) T cell proliferation, CD4^+^ and CD8^+^ T cell activation, and IFN-γ release were determined using ^3^H-thymidine incorporation, flow cytometry and LEGENDplex multiplex assays, respectively (**B**); depletion of LNCaP cells was determined by flow cytometry (**C**). See Supplemental Figure 1A for data from individual patients. (D) PBMC of three patients obtained at the indicated timepoints before (days 1, 22, 43, 64) and after (days 7, 29, 50, 71) treatment were incubated with irradiated LNCaP cells and CC-1 (1 nM). After 72 h, T cell proliferation was assessed by ^3^H-thymidine incorporation. See Supplemental Figure 1C for data from individual patients. (E) Plasma concentrations of CC-1 and soluble interleukin-2 receptor (sIL-2R) were measured at the indicated time points during multiple treatment cycles and serum levels were quantified by ELISA. Exemplary results of one representative patient are shown. See Supplemental Figure 1D for data from two additional patients. (F) Leiden-based clustering of scRNA-sequencing data visualized by UMAP, depicting major immune cell populations in patient PBMC collected at baseline (day 1) and after therapy (day 8), annotated according to canonical lineage markers. UA, unassigned. (G) Dot plots summarizing expression changes of selected genes involved in T cell quiescence/anergy, early activation, and metabolic regulators across time points; dot size indicates the fraction of cells expressing the respective genes; color intensity reflects the mean expression. (H) Gene set enrichment analysis (GSEA) showing significant enrichment of canonical anergy- and hyporesponsiveness-associated transcriptional programs compared to pretreatment samples (day 1). The SAFFORD_T_LYMPHOCYTE_ANERGY pathway (MSigDB) was significantly enriched in day 8 CD4^+^ and CD8^+^ T cells. NES, normalized enrichment score; FDR, false discovery rate. (I) T cell subclustering was visualized by UMAP, resolving distinct functional states including proliferative, effector-like, and hyporesponsive populations that emerge following therapy. (J) Embedding density plots illustrating the distribution of T cell states before treatment (day 1) and after therapy (day 8), documenting the shift toward hyporesponsive and non-proliferative phenotypes. Bar graphs represent the cluster distribution within each condition.

### Format and binder selection for optimized bispecific CD28 costimulation

Monospecific CD28 antibodies require bivalent rather than monovalent binding to exert costimulatory activity (11). Accordingly, bivalent CD28-binding might provide for superior function of costimulatory bsAbs, but comes with the risk of losing target-restricted activity, resulting in side-effects. To address this dichotomy, we generated and characterized four bivalent BiCo formats with irrelevant target-specificity (Figure 2A and Supplemental Figure 3, A and B). Two (bs-CD28-1 and bs-CD28-2) bound CD28 via Fab-arms with accordingly higher affinity to CD28 on T cells, whereas the other two (bs-CD28-3 and bs-CD28-4) contained single-chain anti-CD28 domains fused to heavy or light chains (Supplemental Figure 3C). In combination with submitogenic doses of TCE (CC-1) providing “signal 1”, only bs-CD28-4 displayed strictly target-restricted activity as revealed by the desired lack of T cell stimulation in the absence of TAA-binding (Figure 2B). Accordingly, the format of bs-CD28-4 was selected to generate BiCos directed to endoglin and B7-H3 as targets. These TAAs are broadly expressed on solid tumors and also the tumor-associated extracellular matrix (ECM) including neovasculature (16, 17) and appear thus optimally suitable to bolster CC-1 induced immune responses (Supplemental Figure 3D). From panels of newly generated and previously described (18) mAbs, two endoglin and two B7-H3 binders, respectively, were selected based on optimal affinity (Supplemental Figure 3, E and F). After cloning and characterization of the respective four constructs, two BiCos termed BiCo-1 (endoglin-Kro22xCD28) and BiCo-2 (B7-H3-8H8×CD28) (Supplemental Figure 3, G-I) were selected as lead candidates based on comparative analysis of induction of T cell proliferation (Figure 2C).

**Figure 2.**
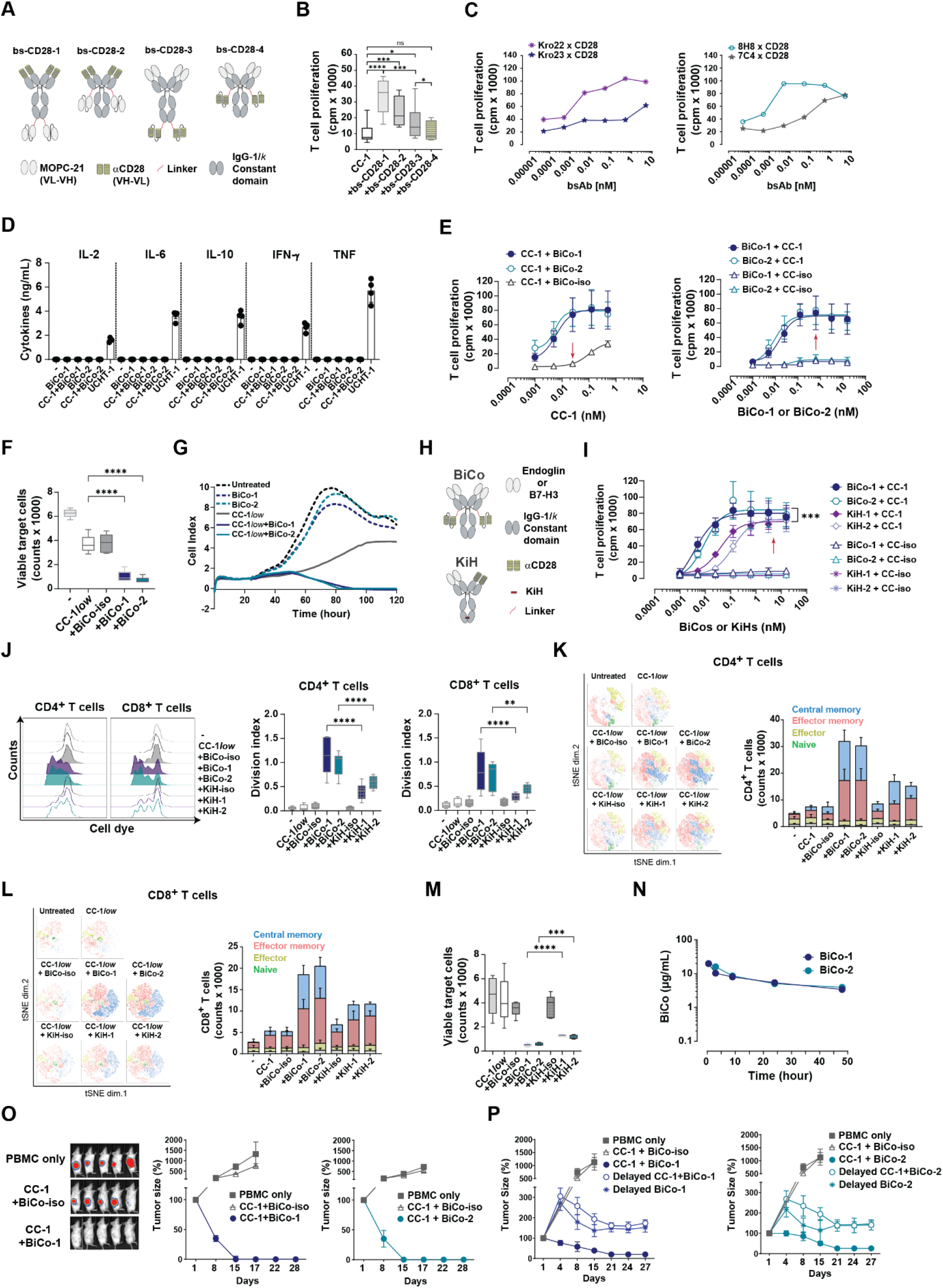
Format selection and functional characterization of BiCOs. (A) Schematic representation of different bispecific CD28 antibody (bs-CD28) formats targeting the irrelevant antigen MOPC-21. The common structure of the different tetravalent formats contains two bivalent binding parts, the first formed by the natural N-terminus of an IgG backbone, the second constituted by two C-terminally fused single chain Fv fragments (scFvs). The CD28 binding part constitutes either the natural N-terminal binding part (variants 1 and 2) or the C-terminal scFv part (variants 3 and 4). The C-terminal scFv parts are either fused to the CH3-(variants 1 and 3) or the CL-domains (variants 2 and 4). The backbone is an Fc silenced IgG1*k* to prevent non-specific Fc-gamma receptor engagement. (B) Irradiated LNCaP cells were cultured with PBMC (n=8) at an E:T ratio of 4:1 in the presence of CC-1 (25 pM) and the indicated bs-CD28 constructs (10 nM). Proliferation was measured after three days by ^3^H-thymidine incorporation. Data are shown as mean ± SD. (C) Irradiated LNCaP-E cells were incubated with PBMCs (n=2 donors) at an E:T ratio of 4:1 in the presence of CC-1 (25 pM) and the indicated concentrations of bs-CD28-4 constructs (Kro22 and Kro23 targeting endoglin; 8H8 and 7C4 targeting B7-H3). After three days, proliferation was assessed using a ^3^H-thymidine incorporation assay. (D) NSG mice (n=4 per group) were injected i.p. with 2×10^7^ human PBMC and i.v. with 20 μg BiCo-1 or BiCo-2 alone or in combination with 0.8 µg CC-1. UCHT1 antibody (anti-CD3) was used as positive control. Serum cytokine levels were measured in blood samples obtained after 24 h by LEGENDplex. (E) PBMC (n=6) were cultured with irradiated LNCaP-E cells and the indicated constructs followed by analysis of proliferation by ^3^H-thymidine incorporation. Left: dose titration of CC-1 in the presence of BiCo-iso, BiCo-1, or BiCo-2 (all at 0.5 nM). Right: dose titration of BiCo-1 and BiCo-2 in the presence of CC-1 or CC-iso (both 25 pM). Arrows indicate concentrations selected for subsequent experiments. Data are shown as mean ± SD. (F) PBMC (n=5 donors) were cultured with LNCaP-E cells at an E:T ratio of 1:1 in the presence of CC-1 (25 pM, CC-1*low*) and the indicated BiCo molecules (0.5 nM). Reduction of target-cells was quantified by flow cytometry after 72 h. (G) Real-time monitoring of LNCaP-E tumor cell growth in the presence of monocyte-depleted PBMC, CC-1 (25 pM, CC-1*low*), and BiCo (0.5 nM) using the xCELLigence system. (H) The BiCo format consists of variant 4 depicted in **A** with a bivalent targeting part directed to endoglin (BiCo-1) or B7-H3 (Bico-2) and a bivalent CD28 binding part (αCD28). The KiH format consists of two natural IgG chains binding the targeting parts (endoglin or B7-H3) and CD28 monovalently. The endoglin-, B7-H3- and CD28-binders used in both formats are identical. (I) PBMC (n=4 donors) were cultured with irradiated LNCaP-E cells in the presence of CC-1 or isotype control (CC-iso) (both at 25 pM) with the indicated concentrations of BiCo-1, BiCo-2, KiH-1, or KiH-2 followed by analysis of proliferation by ^3^H-thymidine incorporation. Data are shown as mean ± SD. Arrows indicate the concentration selected for subsequent experiments. (**J-M**) CellTrace Violet-labeled monocyte-depleted PBMC were cultured with LNCaP-E cells at an E:T ratio of 1:1 with CC-1 (25 pM, CC-1*low*) and BiCo/KiH (both at 5 nM) as indicated. Medium, target cells, and constructs were replenished on day 4, followed by flow cytometric analysis. (**J**) Representative proliferation profiles (left panels) as obtained after 4 days. Division index (n=6 donors) of proliferating CD4^+^ T cells and CD8^+^ T cells was calculated using FlowJo (right panels). (**K-L**) Representative tSNE plots (left) and T cell subset counts (right) for CD4^+^ T cells (**K**) and CD8^+^ T cell (**L**) as determined on day 7 (n=6 donors). (**M**) Tumor cell killing was assessed at day 7 (n=5 donors). (**N**) C57Bl/6 mice (n=4 mice per group) were injected with 20 µg BiCo-1 and BiCo-2 followed by determination of serum concentrations at the indicated time points by Promega assay. (**O-P**) NSG mice bearing established LNCaP-E flank tumors (∼4 mm diameter) were transferred with human PBMCs and treated i.v. with CC-1 (0.8 µg), BiCo-1, BiCo-2, or BiCo-iso (20 µg); Tumor growth was monitored twice weekly. (**O**) Combinatorial treatment with TCE and BiCo was applied on days 1, 8, and 15. Representative bioluminescence images at day 17 (left panel), tumor size as determined by caliper (right panel) are shown. Data represent n=5 mice per group. (**P**) To study effects of delaying BiCo treatment, animals received CC-1 on day 1, followed by BiCo application on day 4 and combined treatment starting at day 8 (delayed BiCo cohorts); alternatively, mice received CC-1 on day 1, followed by combinatorial treatment starting at day 8 (delayed CC-1 + BiCo cohorts). Tumor size was determined by caliper measurement. Data represent n=5 mice per group.

### Functional characterization of BiCo-molecules

Next, in vivo safety of the two BiCo lead candidates was validated in tumor-free humanized mice. In contrast to UCHT1 as control, neither BiCo-1 nor BiCo-2 alone or in combination with CC-1 induced any detectable cytokine release, confirming lack of off-target T cell activation upon BiCo-application and strict target-dependence of the combination regimen (Figure 2D).

Analyses with target-antigen positive versus -negative target cells and dose-response analyses confirmed strictly tumor antigen–dependent activity and demonstrated maximal T cell proliferation at concentrations as low as 25 pM of CC-1 and 0.5 nM BiCo (Figure 2E and Supplemental Figures 4 and 5, A and B). The latter concentrations were used for subsequent experiments. Both BiCos potently enhanced TCE–driven CD4⁺ and CD8⁺ T cell activation, IFN-γ secretion and target killing (Figure 2, F and G; Supplemental Figure 5, C-E). Particularly in long-term assays, combinations of BiCos with low-dose TCE induced significantly enhanced (e.g., for CD8^+^ T cells *P* = 0.03 for BiCo-1 and *P* = 0.004 for BiCo-2) and fully conditional T cell proliferation as well as pronounced expansion of effector- and central-memory subsets as compared to even high doses (1,000 pM) of TCE as single agent (Supplemental Figure 5, F and G).

To benchmark our bivalent BiCo-format against univalent Knob-into-Hole CD28-costimulators (KiH) (19) that are presently being evaluated in clinical trials (NCT06826768, NCT04626635, NCT04590326), we generated KiH molecules comprising the TAA- and CD28-binders that are used in our BiCos (KiH-1/-2) (Figure 2H and Supplemental Figure 6, A and B). Surprisingly, with respect to affinity, the KiH constructs were comparable to our BiCos (Supplemental Figure 3I and Supplemental Figure 6, C-E). Next, we compared the costimulatory potential of our bivalent BiCo format with the monovalent KiH constructs in combination with TCE. The costimulatory activity of our BiCo molecules was clearly superior to that of their KiH counterparts over a broad range of concentrations (Figure 2I). In subsequent analyses, we used 5 nM as saturating concentration for BiCos and KiH molecules and found that our bivalent BiCo format induced significantly superior CD4^+^ and CD8^+^ T cell proliferation (CD4^+^ T cells: *P <* 0.0001 for BiCo versus KiH molecules. CD8^+^ T cells: *P <* 0.0001 for BiCo-1 versus KiH-1 and *P* = 0.0014 for BiCo-2 versus KiH-2), memory formation, and tumor cell killing (*P <* 0.0001 for BiCo-1 versus KiH-1 and *P* = 0.0002 for BiCo-2 versus KiH-2), particularly in long-term assays that are most suitable to unravel the effects of costimulation (Figure 2, J-M and Supplemental Figure 7, A-E). This difference in efficacy could be due to a largely reduced off-rate of the CD28 binding part (by a factor of 5) of our bivalent BiCo molecules (Supplemental Figure 6D), although it cannot be ruled out that the observed difference is due to particular spatial antibody/receptor conformations that are not associated with altered off-rates.

Pharmacokinetic analyses in C57BL/6 mice revealed that the serum half-life of our BiCo format is comparable to that of classical mAbs (Figure 2N; (5)). In NSG mice xenografted with human PBMC, simultaneous combinatorial treatment with CC-1 and BiCos (but not TCE alone) eliminated large established LNCaP-E flank tumors and statistically significantly improved animal survival (*P* <0.0001), as assessed by caliper measurements, bioluminescence imaging and Kaplan-Meyer analysis (Figure 2O, Supplemental Figure 7F). Delayed BiCo application after initial CC-1 single agent application resulted in a less pronounced antitumor effect, but still was by far more effective than single-agent CC-1 treatment, further confirming the potential of CD28-costimulation to reinforce TCE efficacy (Figure 2P).

### BiCos revert and prevent T cell hyporesponsiveness

Analysis of T cell proliferation and tumor cell killing using PBMCs of five patients obtained before (day 1 of cycle 1) and after (day 8) TCE-treatment revealed that application of BiCos completely reverted hyporesponsiveness. Notably, in both, TCE-responsive as well as unresponsive patient T cells, BiCos raised activity levels markedly above those achieved (in responding cells) with a high concentration of TCE alone (Figure 3, A and B, *P* <0.0001). This also remained true when cells obtained during later treatment cycles were analyzed (Figure 3C).

**Figure 3.**
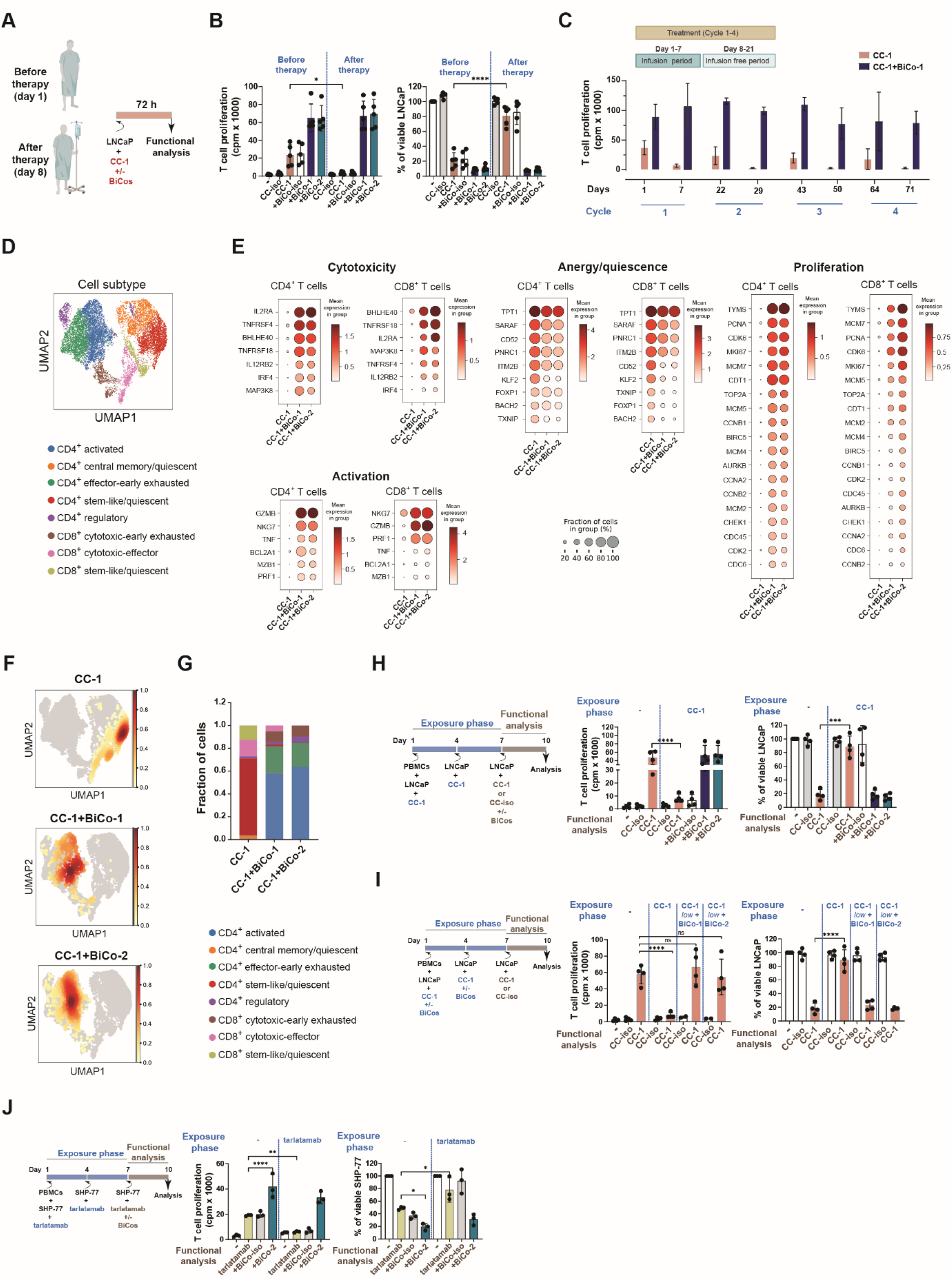
Reversion and prevention of TCE-induced T cell hyporesponsiveness by BiCos. (A) PBMCs were collected from patients before and after TCE treatment with CC-1 (day 1 and day 8, respectively), incubated ex vivo with CC-1 alone or in combination with BiCos, and subsequently subjected to functional analysis. (B) Patient-derived PBMC (n=5) were cultured with LNCaP-E tumor cells at an E:T ratio of 1:1 in the presence of 1 nM CC-1 with or without 0.5 nM of BiCo-1, BiCo-2 or isotype control (BiCo-iso). After 72 h, T cell proliferation was quantified by ^3^H-thymidine incorporation, tumor cell killing was determined by flow cytometry. (C) PBMCs from three patients obtained before and after treatment (day 1 and day 8 of subsequent cycles; see scheme in Figure 1D) were incubated with CC-1 (5 nM) alone or together with BiCo-1 (5 nM). After 72 h, T cell proliferation was assessed by ^3^H-thymidine incorporation. (D) Leiden-based subclustering of scRNA-seq data visualized by UMAP, depicting T cell populations within patient PBMC samples collected after therapy (day 8), and incubated for 3 days in vitro with LNCaP-E and 1 nM CC-1 with or without 0.5 nM BiCos annotated using canonical lineage markers. UMAP visualization of T cell subclustering, resolving distinct functional T cell states. (E) Dot plots summarizing temporal expression changes of selected genes involved in T cell activation, cytotoxicity, proliferation and quiescence. Dot size indicates the fraction of cells expressing the respective gene; and color intensity reflects the mean expression. (**F-G**) Density plots illustrating shifts in patient T cell (day 8) state distributions of T cells treated in vitro with CC-1 alone or CC-1+BiCos (**F**), demonstrating a pronounced transition toward proliferative phenotypes. (**G**) Bar graphs depicting cluster distribution across experimental conditions. (H) PBMCs (n=4) were cultured with LNCaP-E cells at an E:T ratio of 1:2 in the presence of CC-1 (1 nM); medium, target cells, and constructs were replenished on day 4 to mimic chronic exposure. At day 7, PBMCs were re-stimulated with LNCaP-E cells at an E:T ratio of 1:1 in the presence of CC-1 (1 nM) alone or in combination with the indicated BiCo constructs (0.5 nM). On day 10, T cell proliferation was accessed by ^3^H-thymidine incorporation and tumor cell killing was evaluated by flow cytometry. (I) PBMCs (n = 4) were cultured with LNCaP-E cells at an E:T ratio of 1:2 in the presence or absence of CC-1 (1 nM) alone or in combination with BiCo constructs (0.5 nM). Medium, target cells, and constructs were replenished on day 4 and day 7 with (LNCaP-E cells E:T ratio of 1:1 on day 7). At day 10, T cell proliferation was assessed by ^3^H-thymidine incorporation and tumor cell killing was evaluated by flow cytometry. (J) PBMCs (n=3) were cultured with SHP-77 cells at an E:T ratio of 1:2 in the presence of tarlatamab (1 nM). Medium and target cells were replenished on day 4. After the exposure until day 7 mimicking induction of hyporesponsiveness, PBMCs were re-stimulated with SHP-77 cells at an E:T ratio of 1:1 in the presence of tarlatamab (1 nM) or in combination with BiCo-2 (0.5 nM). Proliferation was measured on day 10 by ^3^H-thymidine incorporation, and tumor cell killing was assessed by flow cytometry.

Analysis at the single-cell transcriptional level revealed that BiCo-treatment reinstated a pronounced proliferative cell-cycle program, including upregulation of MCM family members and *CDC* genes alongside enhanced effector programs and activation-associated pathways (*GZMB*, *NKG7*, *TNF*, *IL2RA*, *IRF4*) and reverted hallmark features of T cell anergy/quiescence including downregulation of *KLF2*, *PNRC1*, *TXNIP*, *CD52*, *SARAF*, and *BACH2* (Figure 3, D and E). Additional embedding density analysis (Figure 3, F and G) indicated that BiCo-treatment is associated with CD8⁺ T cells shifting from dysfunctional terminal effector-like states back toward effector-memory populations. In parallel, CD4⁺ T cells transitioned from quiescent/resting phenotypes toward early-activated central memory states, consistent with rebalanced activation and functional recovery.

Induction of hyporesponsiveness upon TCE-treatment was then modelled in vitro by exposing PBMCs and target cells to TCE for 7 days, followed by analysis of responsiveness upon restimulation with the TCE. Alike observed in patients, prolonged exposure to CC-1 resulted in T cell hyporesponsiveness that was dependent on target-engagement, as exposure to CC-1 in the absence of tumor targets did not affect T cell reactivity. Again, in line with the data obtained with patient T cells, hyporesponsiveness could be completely prevented and reverted by presence of BiCos during the exposure to TCE in the first 7 days of culture and during the functional analysis phase, respectively (Figure 3, H and I; Supplemental Figure 7G). Similar results were obtained with tarlatamab (Figure 3J), thereby extending our findings on TCE-induced T cell hyporesponsiveness to other, clinically available formats. Tarlatamab was recently approved for treatment of small cell lung cancer (SCLC)(20) and is a half-life-extended bispecific T cell engager (BiTE) consisting of two single-chain variable fragments (scFvs) that bind DLL3 on tumor cells and CD3 on T cells, fused to a silenced Fc domain.

Together, our results demonstrate that TCE single-agent therapy induces T cell hyporesponsiveness characterized by alterations of intracellular pathways associated with functional anergy/quiescence rather than terminal exhaustion. T cell functionality can be restored and hyporesponsiveness prevented by provision of CD28-signaling using BiCos that reinforce T cell antitumor immunity to an unprecedented level.

## Discussion

TCEs, this is, bsAbs with TAAxCD3-specificity, are well established for treatment of B cell malignancies and currently gain rapid momentum in other tumor entities, exemplified by the recent approval of tarlatamab for treatment of SCLC (7). However, a substantial proportion of patients still does not respond to TCE-treatment, particularly in solid tumors (e.g., 35% objective response rate for tarlatamab (20)). This calls for a better understanding of the mechanisms leading to therapeutic failure as a prerequisite for improving treatment efficacy. When we analyzed T cells of prostate cancer patients undergoing treatment with the PSMAxCD3 TCE CC-1 (12) (NCT04104607), we found that prolonged TCE-treatment had resulted in pronounced functional hyporesponsiveness. The latter was only partly resolved after treatment free intervals as long as two weeks, reflected by a decline of sIL-2R peak serum levels with increasing numbers of treatment cycles. To our knowledge, these data provide the first evidence that TCE-treatment can induce T cell hyporesponsiveness in patients that may impair efficacy against solid tumors. Induction of hyporesponsiveness was also observed upon in vitro stimulation of PBMC with tarlatamab, indicating that the observed phenomenon may generally affect efficacy of TCEs. Future studies should unravel the relationship between T cell hyporesponsiveness and treatment scheme of a given TCE to inform on optimal dosing and efficacy.

Available data document that isolated stimulation of the TCR/CD3 complex causes T cell hyporesponsiveness (2, 21). This state is characterized by impaired IL-2 production, proliferative arrest, and induction of suppressive regulatory programs (3, 22). In line with these findings, we here report for the first time by directly analyzing T cells of patients that exposure to TCE single agent treatment results in suppression of immediate-early AP-1 transcription factors (FOS, JUN family), loss of activation and metabolic fitness programs, and induction of hyporesponsiveness-associated regulators including BACH2, FOXP1, CBLB, PNRC1, and PTPN22, alongside metabolic suppressors such as TXNIP and UCP2 (23, 24). Sustained upregulation of PD-1 or terminal differentiation markers (CD57, KLRG1) was not observed, arguing against irreversible exhaustion or senescence. Instead, the transcriptional and functional features reflect early “paralysis” or costimulation-deficient states described in chronic tumor antigen exposure-models (15). Single-cell analysis further revealed that both CD8⁺ and CD4⁺ T cells stall along differentiation pathways, diverting into dysfunctional effector-like or quiescent-like states. This phenotype is strongly aligned with prior observations that BACH2 antagonizes AP-1–driven effector differentiation, thereby enforcing tolerance-associated or memory-like transcriptional programs when costimulatory signals are limited (23, 24). In tumor and chronic infection settings, elevated BACH2 constrains MYC induction, represses effector lineage commitment, and maintains T cells in a transcriptionally plastic yet functionally restrained quiescent state (14, 25, 26). Together, these findings demonstrate for the first time that TCE single-agent therapy can induce a persistent anergic/quiescent state that restricts durable T cell antitumor immunity and thus efficacy.

We and others have reported previously that costimulatory second signals via the CD28 molecule may counteract T cell hyporesponsiveness mediated by excessive TCR/CD3 stimulation (2, 27, 28). Building on these findings, our early work with bispecific costimulation, and the T cell hyporesponsiveness observed in TCE-treated patients, we reasoned that BiCos may be required to unleash the full potential of bsAb therapy. Concerning the optimization of bispecific costimulation, it is well described that soluble bivalent CD28-antibodies - in contrast to univalent fragments thereof - exert potent costimulatory activity (11). However, this challenges the requirement of target-restriction, the importance of which has been strikingly demonstrated by the Tegenero incident. Conceptually, in the case of CD28-bispecifics, target-restriction of activity usually requires monovalency of the CD28 binding part (29), which in turn limits affinity. We here introduce a tetravalent BiCo format that preserves bivalency for CD28 while still acting in a fully target-restricted manner. Our data indicate that the sterical conformation of the two CD28 scFv-binding moieties allows for agonistic activity only after immobilization upon binding to a target cell. Endoglin (BiCo-1) and B7-H3 (BiCo-2) were chosen as TAAs for lead constructs due to their well characterized expression not only on tumor cells, but also the tumor ECM, which may improve access of immune cells to the tumor site upon therapeutic targeting (16, 17).

In various in vitro models, both BiCo-1 and BiCo-2 potently stimulated T cell anti-tumor reactivity even at very low concentrations of TCE, which holds promise to reduce respective side effects upon clinical application. Combinatorial treatment by far exceeded efficacy of TCEs applied as single agent, even of high concentrations. Superior efficacy of the TCE and BiCo combination was also documented in xenograft mouse models, where only combinatorial treatment achieved tumor elimination. Of note, comparison of our bivalent BiCos to analogous, so far available univalent constructs generated using KiH technology confirmed significantly improved costimulatory capacity of our tetravalent BiCo-format. Somewhat surprisingly, it was not affinity of CD28-binding but rather off-rates that differed significantly and were approx. 5 times faster for univalent constructs, indicating more stable complex formation of BiCos with the CD28 molecule. However, in this context, distinguishing the contribution of kinetic stabilization versus structural receptor clustering is challenging, as the bivalent architecture itself may affect both, receptor geometry and binding kinetics. Thus, the observed efficacy advantage of the BiCo format cannot be unequivocally attributed to either mechanism.

Most importantly, our data show that BiCos can both, reinforce and also restore T cell effector-competence, amongst others by overcoming the hyporesponsive state of T cells of patients treated with TCE as single agent, thereby reinstating proliferation and tumor cell killing. This was mirrored by downregulation of TCE-induced hallmark anergy/quiescence genes and concomitant restoration of activation-, cytotoxicity-, proliferation- and cytokine response-associated gene expression, indicating reversal of “signal 1”-induced hyporesponsiveness to permit progression through G0/G1 arrest toward proliferation and enhanced effector function. These findings align closely with very recent studies showing that CD28 signaling neutralizes anergy by reinforcing IL-2 production, promoting cell-cycle entry, and functionally counteracting BACH2-mediated repression, which diverts both CD8⁺ and CD4⁺ T cells from productive effector fates (14, 26).

In summary, our data document that TCE single-agent treatment can result in T cell hyporesponsiveness that impairs efficacy. Our data obtained in TCE-treated patients are supported by preclinical models with the DLL3xCD3 bsAb tarlatamab, pointing to a class-specific effect of TCE treatment. BiCo administration can preserve proliferative capacity and cytotoxic activity of the cells. Addition of BiCos to hyporesponsive T cells also can restore functionality and revert the TCE-induced molecular signature, confirming that TCE-induced T cell hyporesponsiveness is rather due to an anergic/quiescent state than terminal exhaustion. By elucidating the underlying transcriptional mechanisms and demonstrating their reversibility, we harness conditional CD28-costimulation as pivotal strategy towards achieving durable responses not only with TCEs, but rather as generalizable therapeutic principle for combinations with other “signal-1”-mediating immunotherapies like cancer vaccines. A clinical trial evaluating for the first time a combination of TCEs and BiCos directed to two different TAAs will start enrollment in 2026 and shall provide clinical evidence for our concept to increase both safety and efficacy of T cell recruiting bsAb, holding promise to bring therapy with these reagents to an unprecedented level.

## Methods

### Sex as a biological variable

All patients included in this study were male and treated for metastatic castration-resistant prostate cancer. In vivo experiments were performed using female mice, as this allows for more homogeneous tumor engraftment and reduces variability across experimental groups.

### Patient samples and cells

PBMC samples of five patients with metastatic castration-resistant prostate cancer treated within a phase I clinical trial (ClinicalTrials.gov identifier: NCT04104607) evaluating the PSMAxCD3 bsAb CC-1 (12) were obtained prior to and after 7 days of continuous application of CC-1: 78 µg on day 1, 215 µg on day 2 and 826 µg daily from day 3 to 7 (30). PBMCs of patients and healthy donors were isolated from heparinized peripheral blood by density-gradient centrifugation using Biocoll separating solution (Biochrom, Berlin, Germany).

The human prostate cancer cell line LNCaP was obtained from the Deutsche Sammlung von Mikroorganismen und Zellkulturen (DSMZ) and SHP-77 cell line from American Type Culture Collection (ATCC). To generate endoglin-expressing (LNCaP-E) and B7-H3–deficient (LNCaP^KO^) cells, parental LNCaP cells were transduced with lentiviral vectors encoding firefly luciferase and human endoglin and lentiviral vectors targeting B7-H3, respectively, as previously described (18). LNCaP-E and LNCaP^KO^ cells were cultured under puromycin selection (10 µg/mL), with LNCaP^KO^ additionally maintained in Geneticin (1 µg/mL). All cells were routinely tested for mycoplasma contamination.

### Generation of mAbs to endoglin

Eight-week-old BALB/c mice were immunized intraperitoneally with 5×10⁶ Sp2/0-Ag14 cells transfected with endoglin. Up to four immunizations were administered at two-week intervals, followed by a final boost four days prior to fusion. Splenocytes were harvested and fused with Sp2/0-Ag14 myeloma cells using polyethylene glycol (PEG 1500). Fused cells were cultured in IMDM supplemented with fetal- and horse-serum and distributed into 96-well plates. Hybridomas were selected using HAT medium and cultured for 7–10 days. Supernatants were screened by flow cytometry for antibody production and antigen specificity. Positive clones were re-tested, subcloned by limiting dilution, validated for monoclonality and cryopreserved.

### Generation and purification of bispecific antibodies

Variable domains specific for endoglin, B7-H3, and MOPC-21 (isotype control) were codon-optimized for expression in CHO cells using the GeneArt GeneOptimizer tool (Thermo Fisher Scientific). VH, VL, and scFv sequences were synthesized *de novo* by GeneArt. The variable domains were cloned into a human IgG1κ-based IgGsc backbone containing LALA mutations (L234A/L235A) and YTE mutations (M252Y/S254T/T256E) for Fc-silencing and extended serum half-life. The CD28 scFv derived from clone 9.3 was covalently linked to the C-terminus of the light- or heavy chain via a flexible glycine-serine linker. For KiH bispecific constructs, heterodimerization-promoting mutations were introduced into the CH3 domain of the Fc region (knob: T366W; hole: T366S/L368A/Y407V) as previously reported (31). BsAbs were produced in CHO cells, purified by protein A affinity chromatography, and biochemically characterized by sodium dodecyl sulfate–polyacrylamide gel electrophoresis (SDS-PAGE) and size-exclusion chromatography as previously described (18). Endotoxin levels in all bsAbs were <0.5 U/mL as determined by Limulus amebocyte lysate assay (Endosafe®, Charles River).

### Binding kinetics and affinity measurements

Antibody binding kinetics and affinities were determined by surface plasmon resonance (SPR) and biolayer interferometry (BLI). For SPR analyses, mAbs specific for endoglin or B7-H3 were captured on Protein A-coated sensor chips (GE Healthcare) using a Biacore X instrument. Recombinant His-tagged endoglin or B7-H3 protein produced in CHO cells was injected as analyte at increasing concentrations. Association and dissociation phases were recorded, and kinetic parameters (*k_on_, k_off_*) and equilibrium dissociation constants (*K_D_*) were calculated using BIAevaluation software v4.1 (Cytiva). BLI analyses were conducted by Sino Biological, Inc. Binding affinity was assessed by biolayer interferometry using Octet RED96 (Sartorius). Protein A biosensors were loaded with the mAbs or bsAbs and incubated with serial dilutions of recombinant endoglin, B7-H3 or CD28. Association and dissociation were monitored under constant agitation. Data were processed and fitted using Octet Data Analysis software (v11.1) to derive kinetic and affinity constants.

Binding of bsAbs to antigens expressed on tumor cells and CD28 on T cells was determined using LNCaP-E cells and PBMCs, respectively. Primary antibodies were detected using PE-conjugated goat anti-human F(ab)2 fragments (Jackson ImmunoResearch, West Grove, PA, USA). Flow cytometric analysis was performed using BD FACSCantoTM II and BD FACSCaliburTM (BD Biosciences, San Jose, CA, USA) and data were analyzed using FlowJo (FlowJo LLC, Ashland, Oregon, USA). EC_50_ values were calculated using GraphPad Prism9 (GraphPad Software).

### Single-cell transcriptomic profiling and computational analysis

Sample preparation of PBMCs from prostate cancer patients was performed according to the 10× Genomics Feature Barcode protocol for cell surface protein labeling in scRNAy-seq (CG000149). To enable sample multiplexing and limit batch effects, cells from individual donors were labeled with TotalSeq-C hashtag oligonucleotide–conjugated antibodies (Supplemental Table 1). Subsequently, single cells were encapsulated into Gel Beads-in-Emulsion (GEMs) together with barcoded gel beads and reverse transcription reagents using the Chromium X instrument (10× Genomics). Subsequent library preparation included single-cell 5′ gene expression (GEX), T cell receptor (TCR) variable-diversity-joining (VDJ), and cell surface protein (SP) libraries generated using Chromium Next GEM Single Cell 5′ HT Reagent Kit v2 (Dual Index) (CG000424; 10× Genomics, PN-1000356), Chromium Library Construction Kit (PN-1000352), Chromium Single Cell Human TCR Amplification Kit (PN-1000252), and Chromium 5′ Feature Barcode Kit (PN-1000541) according to manufacturer’s protocols. The resulting libraries were pooled and subjected to high-throughput sequencing using Illumina NovaSeq 6000 systems. Barcode processing, alignment, VDJ annotation, and single-cell 5′ gene counting were performed using Cell Ranger Software version 7.1.0 (10X Genomics) aligned to the GRCh38-202-A reference genome. Further data processing, visualization, and analysis were performed using scanpy version 1.11.1 for all samples (32). Demultiplexing was further handled with R library Seurat (33, 34). Genes below a threshold of 3 regarding the number of cells expressing the gene and the number of unique UMI counts as well as mitochondrial genes, ribosomal genes, HG, lncRNA and pseudogenes were excluded from clustering. Neighborhood graphs and UMAP embedding were computed using UMAP algorithm, and unsupervised clustering was performed using Leiden algorithm. Unsupervised clustering was first conducted on the entire dataset to identify major immune cell populations. When focusing on T cell subsets, B cells and unidentified populations were excluded from analysis.

Parameters used for clustering of the ex vivo and in vitro samples were n_pcs = 25, n_neigbors = 30, min_dist = 0.5, resolution = 0.21 and n_pcs = 13, n_neigbors = 25, min_dist = 0.3, resolution = 0.63, respectively. CD3^+^ clusters were selected for sub-clustering using the same algorithms with parameters for in vivo and in vitro samples being n_pcs = 5, n_neigbors = 5, min_dist = 0.7, resolution = 0.1 and n_pcs = 31, n_neigbors = 15, min_dist = 0.45, resolution = 0.4, respectively. DEGs were calculated using Wilcoxon rank-sum test with Benjamini-Hochberg adjustment and tie correction. Gene-Set-enrichment analysis (GSEA) was conducted using the GSEApy prerank function (35) (version 1.1.11) to evaluate functional changes between time points day 8 and day 1 in CD4^+^ and CD8^+^ T cells using Safford_T_Lymphocyte_Anergie Pathway from MSigDB C2 (36) and GSE46242_TH1_VS_ANERGIC_TH1_CD4_TCELL_UP from ImmuneSigDB (37). The scores served as rank metric, and the GSEApy function prerank was run with 2000 permutations (permutation_num=2000). For both cell types, the enrichment (ES) and normalized enrichment score (NES), the nominal *p*-value and Benjamini-Hochberg adjusted q-value (FDR) were calculated. An adjusted *p*-value < 0.05 was set as statistical significance cutoff. The leading-edge genes that contributed most to the signal and are up-regulated at day 8 were extracted and listed separately in Supplemental Table 1.

### T cell proliferation

PBMCs depleted of monocytes were cultured with target cells with or without bsAbs. After 48 h, cells were pulsed with ^3^H-methyl-thymidine (0.5 μCi/well) for 20 h followed by determination of incorporated radioactivity liquid scintillation counting in a 2450 Microplate counter (Perkin Elmer). Alternatively, PBMCs were labeled with 2 µM CellTrace Violet (Thermo Fisher Scientific) and cultured with target cells in the presence or absence of bsAbs. On day 3, tumor cells and bsAbs were replenished, and T cell proliferation was determined after a total of 4 or 6 days by flow cytometry. T cell subsets were analyzed for CD4-APC/Cy7, CD8-FITC, CD45RO-PE/Cy7, and CD62L-Pacific Blue (BioLegend, San Diego, CA, USA) expression by flow cytometry.

### Cytotoxicity assays

To determine target cell lysis using flow cytometry-based assays, monocyte-depleted PBMCs were cultured with tumor cells in the presence or absence of the indicated bsAbs for 24 or 72 hours. T cells were identified using CD4-Pacific Blue and CD8-FITC, activation status was determined by CD69-APC/Cy7 and CD25-PE, tumor cells were identified using EpCAM-PE/Cy7 and dead cells were excluded using 7-AAD (BioLegend, San Diego, CA, USA). Calibration beads (Sigma-Aldrich, St. Louis, MO, USA) were employed to control for sample volume and to adjust for the number of viable target cells. Antibodies are listed in Supplemental Table 2. For real-time cytotoxicity measurements, the xCELLigence RTCA system (Roche Applied Science, Penzberg, Germany) was used to monitor tumor cell killing every 30 minutes over a period of 120 hours.

### Cytokine secretion

PBMCs were incubated with tumor cells in the presence of CC-1 and the indicated bsAbs. After 24 h, supernatants were collected and cytokine levels (IL-2, IFN-γ) were measured using Legendplex assays (BioLegend, San Diego, CA, USA) according to manufacturer’s protocol. For patient, soluble IL-2 receptor (sIL-2R), blood samples were collected in Li-heparin monovettes and plasma was obtained by centrifugation. sIL-2R concentrations were determined from heparin plasma in the central laboratory using a commercial ELISA kit (IBL, Germany) according to the manufacturer’s instructions.

### Analysis of T cell hyporesponsiveness in vitro

To evaluate T cell hyporesponsiveness in vitro, PBMCs of healthy human donors were cultured with tumor cells (E:T = 2:1) with or without TCE (1 nM) alone or in combination with BiCos (0.5 nM). Culture medium, tumor cells, and antibodies were replenished on day 4. At day 7, PBMCs were again stimulated with tumor target cells (E:T = 1:1) and TCEs with and without BiCos. At day 10, T cell proliferation was assessed by ^3^H-thymidine incorporation, and CD8⁺ T cell activation and tumor cell killing were evaluated by flow cytometry. Flow cytometric data were acquired and analyzed as described above.

Surface marker expression of patient PBMCs depicted in the Supplemental Figure 1, C and D, was determined by flow cytometry without prior in vitro stimulation. Antibodies are listed in Supplemental Table 2, and samples were acquired on a Cytek Aurora spectral cytometer with automated compensation calculation.

### In vivo studies

All animal experiments were conducted as described previously (12, 18). All mouse strains were bred and maintained under specific-pathogen-free (SPF) conditions. Age- and sex-matched mice were used for all in vivo studies, and animals were monitored daily to assess health status. Humane endpoints were predefined, and mice meeting these criteria were euthanized accordingly. Blinding was not implemented in this study.

For assessment of pharmacokinetics: 20 µg of the indicated constructs were intravenously injected into C57BL/6 mice and serum concentrations were determined using bioluminescent T cell-based assays (Promega, Madison, USA).

For determination of anti-tumor activity, female NSG mice were subcutaneously injected in the right flank with LNCaP-E cells transduced with firefly luciferase to establish tumors of approximately 5 mm in diameter. On days 1, 8, and 15, mice received intravenous injections of 1×10⁷ human PBMCs with or without CC-1 (0.8 µg) in combination with BiCo-1, BiCo-2 or isotype control (20 µg). In delayed BiCo cohorts, mice received CC-1 alone on day 1, followed by BiCo-1 or BiCo-2 on day 4 and subsequent CC-1/BiCo combination treatment on days 8 and 15. In delayed combination cohorts, mice received CC-1 alone on day 1, followed by initiation of CC-1 plus BiCo treatment on day 8 and continuation on day 15. Tumor growth was monitored twice weekly, and mice were euthanized when tumors reached 15 mm in diameter. Bioluminescence images were acquired on an IVIS Lumina II system (PerkinElmer) and quantified using Living Image software.

To evaluate toxicity, NSG mice received intravenous injections of 20×10⁶ human PBMCs with or without 20 µg of the indicated antibodies. Levels of serum cytokine levels were measured 24 hours later using the LEGENDplex™ Human Th1 Cytokine Panel (BioLegend) according to the manufacturer’s instructions.

### Statistical Analysis

Analyses were performed using GraphPad Prism software. Data are presented as mean ± standard deviation (SD) or as individual values. Comparisons between groups were conducted using, two-way ANOVA, one-way ANOVA, Student’s t-test, Wilcoxon signed-rank test, or the nonparametric Mann–Whitney test. Normality was assessed for each group using the Shapiro–Wilk test. Homogeneity of variance was assessed using the Brown–Forsythe test.

### Study approval

Human PBMC samples from patients with metastatic castration-resistant prostate cancer enrolled in a phase I clinical trial (NCT04104607) and from healthy donors were obtained with written informed consent in accordance with the Declaration of Helsinki. Animal experiments were conducted in accordance with the principles of replacement, reduction, and refinement (“3Rs”), following the ARRIVE guidelines and European animal protection laws (Directive 2010/63/EU). Experimental protocols were approved by the Institutional Animal Care and Use Committee of the University of Tübingen (M02/22 G), in compliance with German federal and state regulations.

## Supporting information

Zekri et al. Supplemental material

## Data availability

All raw data associated with the main article and supplemental material are included in the Supporting Data Values files. Single-cell transcriptomic data are available in the Gene Expression Omnibus (GEO) database (GSE333581). Other data are available upon request.

## Author contributions

**L. Zekri:** Conceptualization, resources, data curation, formal analysis, supervision, validation, investigation, visualization, methodology, writing–original draft, project administration, writing–review and editing. **N. Köhler:** Data curation, software, formal analysis, supervision, funding acquisition, investigation, methodology, project administration, writing–review and editing. **A. Metzger:** Data curation, software, formal analysis, validation, methodology. **N. Prakash:** Data curation, formal analysis, validation. **J.S. Heitmann:** Resources. **M. Engel:** Data curation, formal analysis, validation. **T. Manz:** Data curation, formal analysis, validation, investigation. **S. Müller:** Data curation, formal analysis, validation. **S. Hörner:** Data curation, formal analysis, validation. **K. Schwartz:** Resources. **M. Zwick:** Data curation, formal analysis, validation. **I. Hagelstein:** Data curation, formal analysis, validation. **N. Brückner:** Data curation, formal analysis, validation. **M. Pflügler:** Resources. **J. Leibold:** formal analysis, validation. **M. Börries:** Resources, software, supervision, funding acquisition, validation, writing–review and editing. G. Jung: Conceptualization, resources, supervision, funding acquisition, validation, investigation, visualization, methodology, project administration, writing–review and editing. **H. S. Salih:** Conceptualization, resources, supervision, funding acquisition, validation, investigation, visualization, methodology, project administration, writing–review and editing.

## Acknowledgements

The authors thank A. Dobler and C. Walker for excellent technical assistance, M. Märklin for support with mouse work, and A. Rausch for support with patient PBMCs. Flow cytometry sample acquisition was performed at the Flow Cytometry Core Facility Tübingen.

