## Supplementary material for "Bivalent bispecific CD28-antibodies reinforce T-cell responsiveness and revert anergy/quiescence in patients treated with bispecific CD3-antibodies": Zekri et al. Supplemental material

### Shared last authors

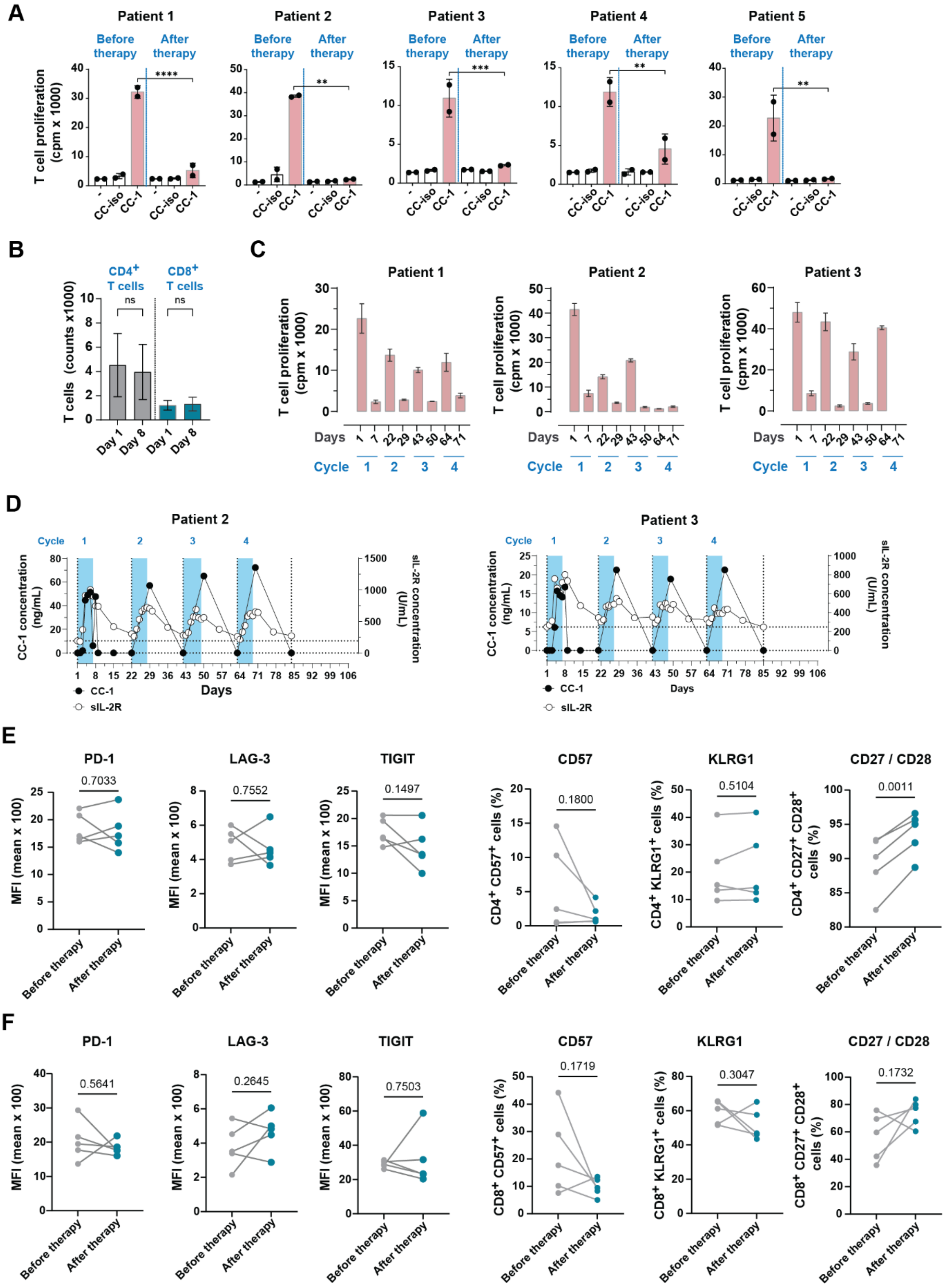
A
Patient 1
Patient 2
Patient 3
Patient 4
Patient 5
Before therapy
After therapy
T cell proliferation (cpm x 1000)
-
CC-iso
CC-1
****
**
***
B
CD4+ T cells
CD8+ T cells
ns
T cells (counts x1000)
Day 1
Day 8
C
Days
Cycle
D
CC-1 concentration (ng/mL)
sIL-2R concentration (U/mL)
CC-1
sIL-2R
E
PD-1
LAG-3
TIGIT
CD57
KLRG1
CD27 / CD28
MFI (mean x 100)
CD4+ CD57+ cells (%)
CD4+ KLRG1+ cells (%)
CD4+ CD27+ CD28+ cells (%)
0.7033
0.7552
0.1497
0.1800
0.5104
0.0011
F
CD8+ CD57+ cells (%)
CD8+ KLRG1+ cells (%)
CD8+ CD27+ CD28+ cells (%)
0.5641
0.2645
0.7503
0.1719
0.3047
0.1732

**Supplemental Figure 1 | Induction of hyporeponsiveness and surface marker expression on T cells from TCE treated patients**

(**A**) PBMC of five mCRPC patients treated with the TCE obtained prior to and after 7 days of therapy were cultured with LNCaP-E tumor cells at an E:T ratio of 1:1 with or without 1 nM CC-1 or isotype control (CC-iso). After 72 h, proliferation was measured by $^{3}$H-thymidine incorporation assay (A, cpm, counts per minute).

(**B**) CD4$^{+}$ and CD8$^{+}$ T cell counts in the patient PBMC samples collected before (day 1) and after treatment (day 8) used in figure 1B and 3B (n = 5) were determined by flow cytometry. Data represent mean ± SEM.

(**C**) PBMCs from three patients were obtained at the indicated time points before treatment (days 1, 22, 43, and 64) and after treatment (days 7, 29, 50, and 71) and incubated with irradiated LNCaP cells and CC-1 (1 nM). After 72 hours, T cell proliferation was assessed by $^{3}$H-thymidine incorporation. Of note, for day 71 of patient 3, no sample was available for analysis. Data represent results of the individual patients of which combined results were shown in Figure 1D.

(**D**) Plasma concentrations of CC-1 and soluble interleukin-2 receptor (sIL-2R) were measured in 2 patients. Samples were collected at the indicated time points during treatment, and serum concentrations of sIL-2R were quantified using ELISA.

(**E-F**) Patient PBMC samples shown in Supplemental Figure 1A, were analyzed for the expression of the indicated markers of T cell exhaustion, anergy, and senescence by flow cytometry on CD4$^{+}$ (**E**) and CD8$^{+}$ (**F**) T cells.

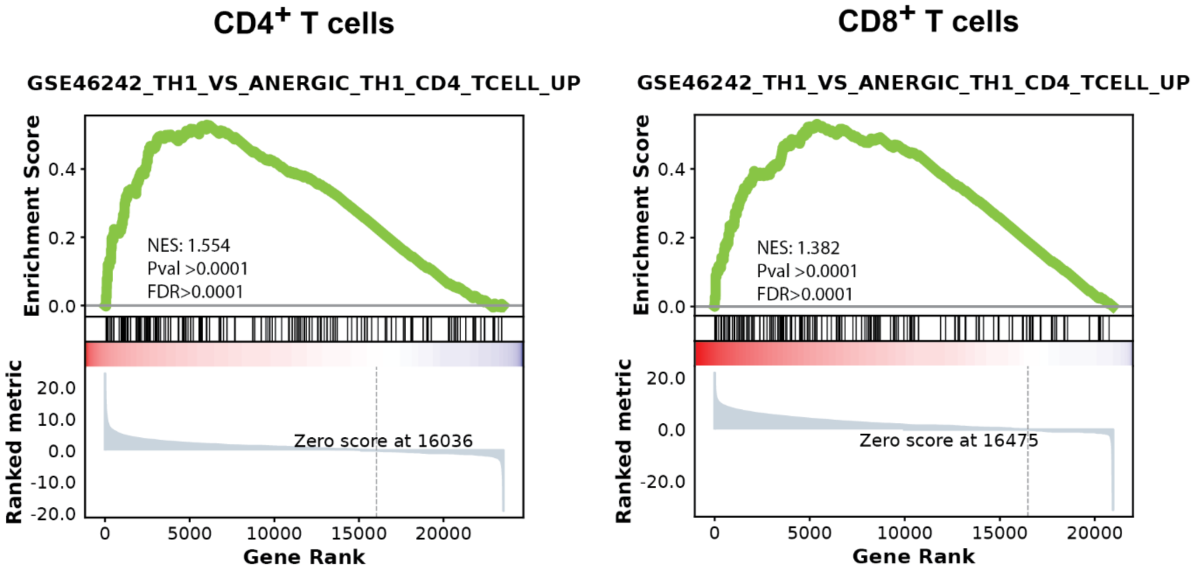


**Supplemental Figure 2 | Induction of anergy-associated transcriptional programs in T cells following TCE treatment**

Gene set enrichment analysis (GSEA) demonstrating significant enrichment of canonical anergy- and hyporesponsiveness-associated transcriptional programs in post-treatment samples compared with pretreatment (day 1). The ImmuneSigDB TH1_vs_Anergic TH pathway was significantly enriched in both $CD4^+$ and $CD8^+$ T-cells at day 8. NES, normalized enrichment score; FDR, false discovery rate.

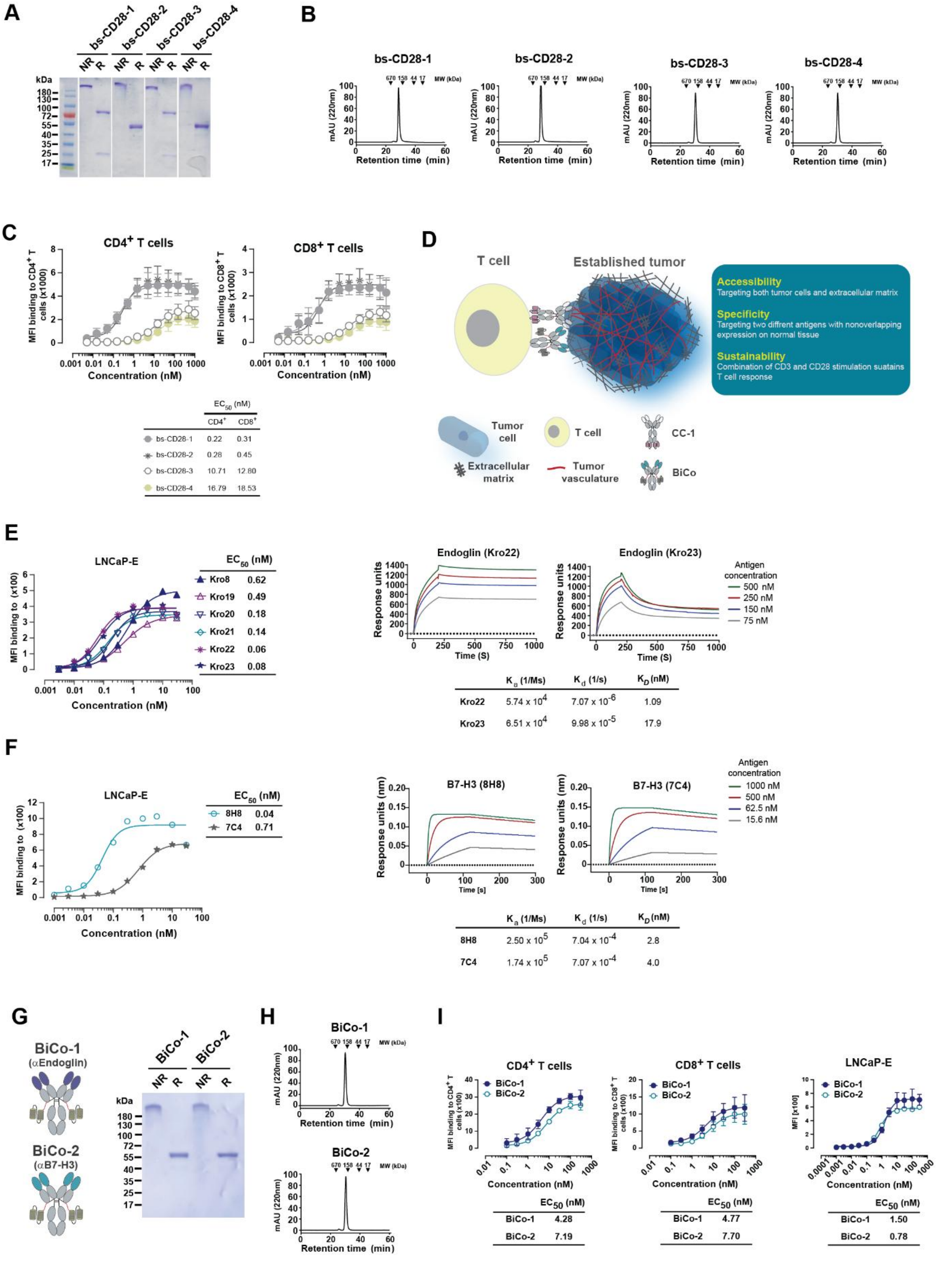
A
bs-CD28-1
bs-CD28-2
bs-CD28-3
bs-CD28-4
NR R
kDa
180
130
100
72
55
40
35
25
17
B
bs-CD28-1
bs-CD28-2
bs-CD28-3
bs-CD28-4
670 158 44 17 MW (kDa)
mAU (220nm)
Retention time (min)
C
CD4+ T cells
CD8+ T cells
MFI binding to CD4+ T cells (x1000)
MFI binding to CD8+ T cells (x1000)
Concentration (nM)
EC50 (nM)
CD4+ CD8+
bs-CD28-1 0.22 0.31
bs-CD28-2 0.28 0.45
bs-CD28-3 10.71 12.80
bs-CD28-4 16.79 18.53
D
T cell
Established tumor
Accessibility
Targeting both tumor cells and extracellular matrix
Specificity
Targeting two diffrent antigens with nonoverlapping expression on normal tissue
Sustainability
Combination of CD3 and CD28 stimulation suatains T cell response
Tumor cell
T cell
CC-1
Extracellular matrix
Tumor vasculature
BiCo
E
LNCaP-E
MFI binding to (x100)
Concentration (nM)
EC50 (nM)
Kro8 0.62
Kro19 0.49
Kro20 0.18
Kro21 0.14
Kro22 0.06
Kro23 0.08
Endoglin (Kro22)
Endoglin (Kro23)
Response units
Time (S)
Antigen concentration
500 nM
250 nM
150 nM
75 nM
Ka (1/Ms) Kd (1/s) KD (nM)
Kro22 5.74 x 10^4 7.07 x 10^-6 1.09
Kro23 6.51 x 10^4 9.98 x 10^-5 17.9
F
LNCaP-E
MFI binding to (x100)
Concentration (nM)
EC50 (nM)
8H8 0.04
7C4 0.71
B7-H3 (8H8)
B7-H3 (7C4)
Response units (nm)
Time [s]
Antigen concentration
1000 nM
500 nM
62.5 nM
15.6 nM
Ka (1/Ms) Kd (1/s) KD (nM)
8H8 2.50 x 10^5 7.04 x 10^-4 2.8
7C4 1.74 x 10^5 7.07 x 10^-4 4.0
G
BiCo-1 (αEndoglin)
BiCo-2 (αB7-H3)
BiCo-1
BiCo-2
NR R
kDa
H
BiCo-1
BiCo-2
670 158 44 17 MW (kDa)
mAU (220nm)
Retention time (min)
I
CD4+ T cells
CD8+ T cells
LNCaP-E
BiCo-1
BiCo-2
MFI binding to CD4+ T cells (x100)
MFI binding to CD8+ T cells (x100)
MFI [x100]
Concentration (nM)
EC50 (nM)
BiCo-1 4.28
BiCo-2 7.19
EC50 (nM)
BiCo-1 4.77
BiCo-2 7.70
EC50 (nM)
BiCo-1 1.50
BiCo-2 0.78

**Supplemental Figure 3 | Selection of format and target binders for generation of BiCo molecules**

(**A**) SDS-PAGE of different CD28 bispecific antibody (bs-CD28) formats targeting non-related antigen (MOPC-21). NR: non-reduced; R: reduced.

(**B**) Size exclusion chromatography of bs-CD28 molecules using Superdex S200 Increase 10/300GL column.

(**C**) Binding of the indicated bs-CD28 constructs to CD28 expressing $CD4^{+}$ and $CD8^{+}$ T-cells within PBMC (n=4 donors). Data are shown as mean ± SD. $EC_{50}$ values were calculated using GraphPad Prism software.

(**D**) Schematic illustration of combinatorial approach: BiCos are directed to endoglin or B7-H3 that are expressed on tumor cells and tumor-associated extracellular matrix/neovasculature. BiCos are combined with TCEs directed to PSMA, which is co-expressed with endoglin or - B7-H3 exclusively in tumor- but not normal tissue. Since TCEs and BiCos are functionally interdependent, sustained T cell activation requires the presence of both antigens at the tumor site, resulting in largely increased specificity of the combinatorial approach.

(**E**) Binding of newly generated anti-endoglin mAbs to LNCaP-E cells (left panel). Surface plasmon resonance (SPR) sensorgrams (right panels) using a Biacore X instrument at 25°C. Kinetic parameters derived from SPR analysis are summarized in the accompanying table.

(**F**) Binding of two anti–B7-H3 mAbs, 7C4 and 8H8, to LNCaP-E cells as determined by flow cytometry (left panel). Biolayer interferometry (BLI; Octet) analysis (right panels) of interactions between recombinant human B7-H3 and the two mAbs. Representative association and dissociation sensorgrams are shown for the indicated analyte concentrations.

(**G**) Schematic illustration and SDS-PAGE analysis of BiCo-1 (endoglinxCD28, Kro22x9.3) and BiCo-2 (B7-H3xCD28, 8H8x9.3); NR: non-reduced; R: reduced.

(**H**) Size exclusion chromatography of BiCo-1 and BiCo-2 using Superdex S200 Increase 10/300GL column.

(**I**) Binding of the indicated BiCo constructs to CD28 expressing $CD4^{+}$ and $CD8^{+}$ T-cells (n=4 donors) and to LNCaP-E cells was assessed by flow cytometry. Data are shown as mean ± SD. $EC_{50}$ values were calculated using GraphPad Prism software.

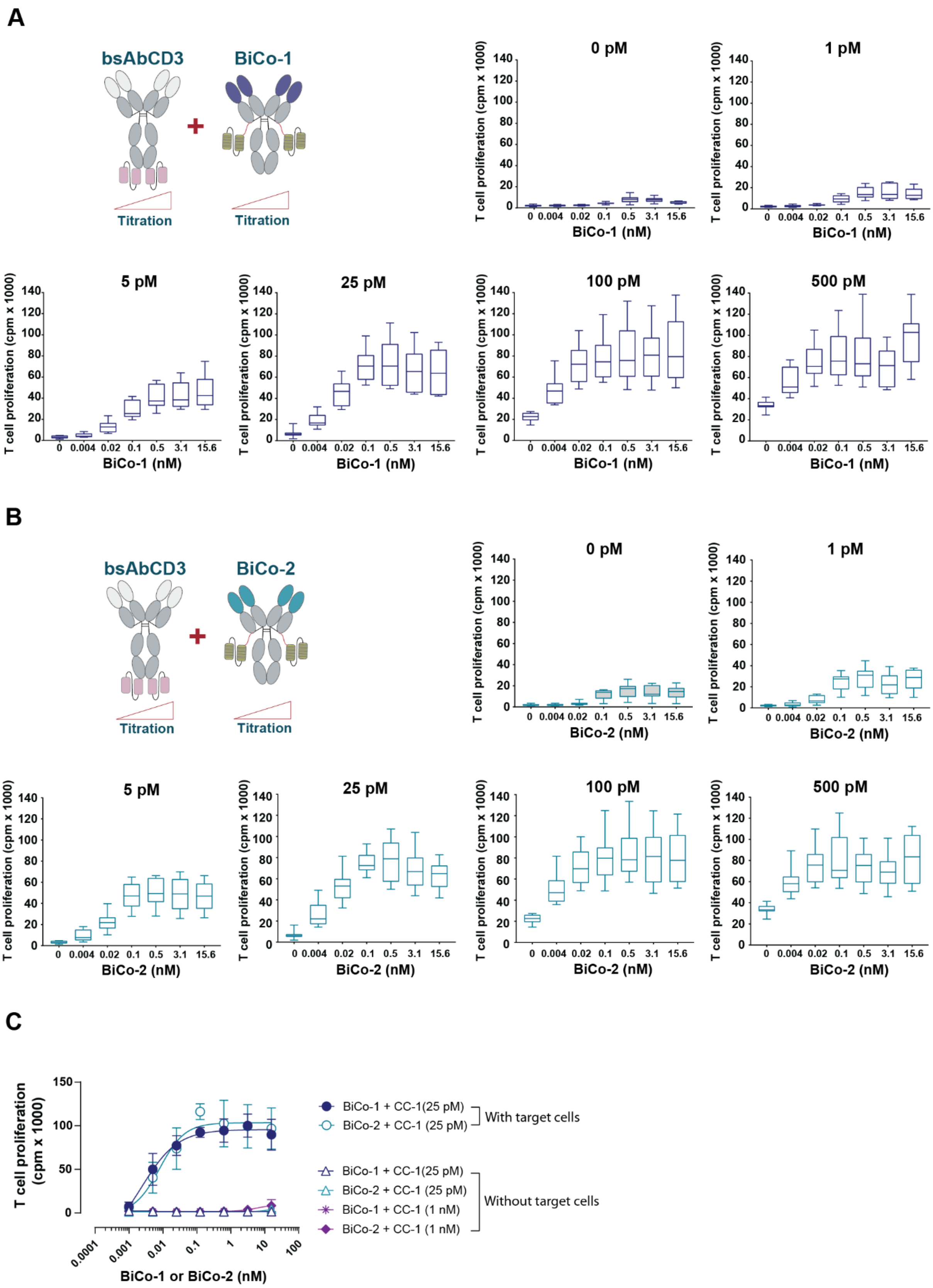
A
bsAbCD3
BiCo-1
Titration
Titration
0 pM
1 pM
5 pM
25 pM
100 pM
500 pM
T cell proliferation (cpm x 1000)
BiCo-1 (nM)
0 0.004 0.02 0.1 0.5 3.1 15.6
B
bsAbCD3
BiCo-2
Titration
Titration
0 pM
1 pM
5 pM
25 pM
100 pM
500 pM
T cell proliferation (cpm x 1000)
BiCo-2 (nM)
C
T cell proliferation (cpm x 1000)
BiCo-1 or BiCo-2 (nM)
BiCo-1 + CC-1(25 pM)
BiCo-2 + CC-1 (25 pM)
With target cells
BiCo-1 + CC-1(25 pM)
BiCo-2 + CC-1 (25 pM)
BiCo-1 + CC-1 (1 nM)
BiCo-2 + CC-1 (1 nM)
Without target cells

**Supplemental Figure 4 | BiCos mediate potent co-stimulation over a wide range of TCE concentrations**

(**A-B**) Proliferation of T cells cultured with irradiated LNCaP-E target cells in the presence of the TCE (CC-1) and BiCo-1 (**A**) or BiCo-2 (**B**) at the indicated concentrations. Data are shown as means ± SD using PBMC of six Independent donors.

(**C**) The effect of the indicated concentrations of BiCo-1 or BiCo-2 on T cell proliferation (n=3) in the presence of two concentrations of CC-1 (25 pM or 1 nM) was determined in cultures with or without irradiated LNCaP-E cells. Data are presented as mean ± SD of experiments with PBMCs from three independent donors.

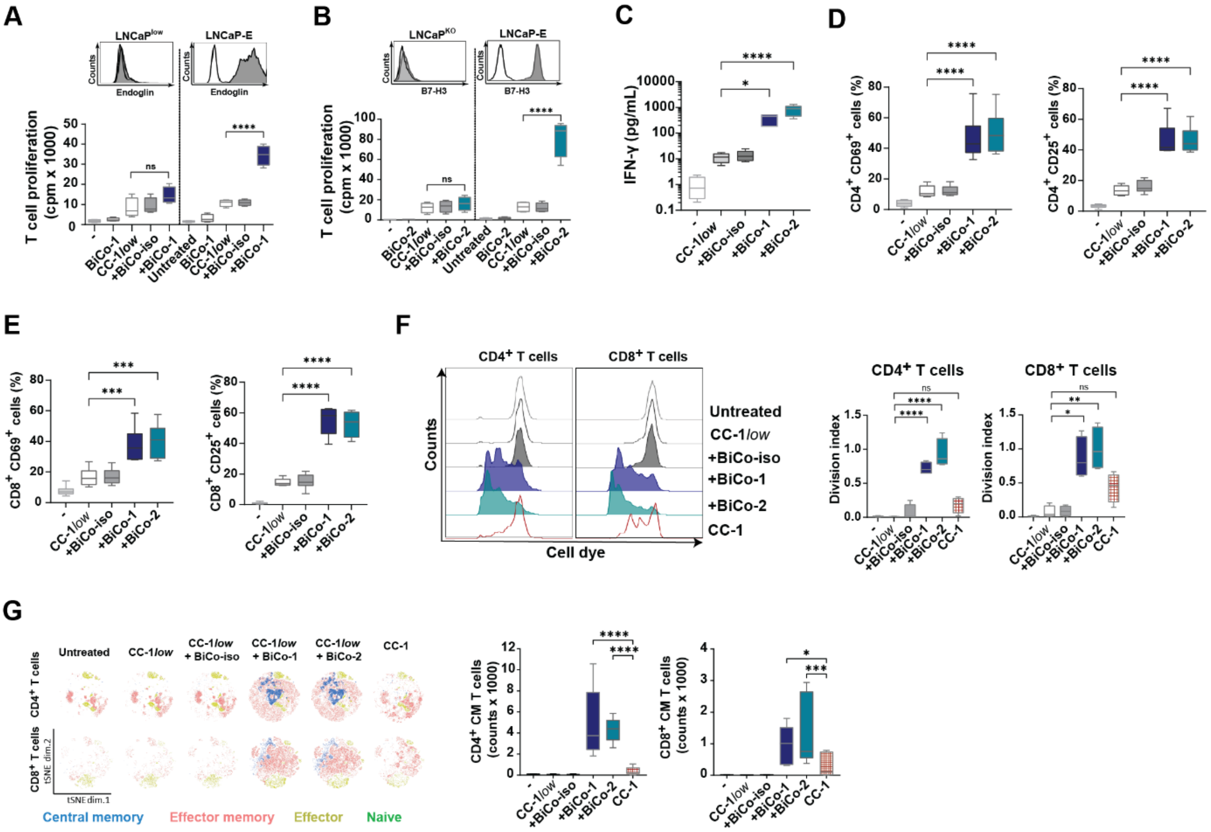


#### Supplemental Figure 5 | Target-restricted CD28 costimulation enhances TCE–mediated T-cell function

(**A-B**) LNCaP$^{low}$ and LNCaP-E cells expressing low and high levels, respectively, of endoglin (**A**), and LNCaP$^{KO}$ and LNCaP-E cells expressing no and high levels, respectively, of B7-H3 (**B**) were analyzed for target expression by flow cytometry (upper panels). PBMCs (n=4 donors) were cultured with the indicated irradiated LNCaP cells at an E:T ratio of 4:1 in the presence of CC-1 (25 pM) with or without BiCos (0.5 nM). Proliferation was measured after three days by $^{3}$H-thymidine incorporation.

(**C-E**) PBMCs (n=5 donors) were cultured with LNCaP-E cells at an E:T ratio of 1:1 in the presence of CC-1 (25 pM, CC-1*low*) and BiCo molecules (0.5 nM), IFN-γ levels in culture supernatants were measured after 24 h incubation using LEGENDplex (**C**). Expression of T cell activation markers CD69 and CD25 on CD4$^{+}$ (**D**) and CD8$^{+}$ T cells (**E**) by flow cytometry after 72 h.

(**F-G**) CellTrace Violet-labeled PBMC were cultured with LNCaP-E cells at an E:T ratio of 1:1 in the presence or absence of the indicated constructs. CC-1 was used at 1 nM or 25 pM (low), alone or in combination with the indicated BiCos (0.5 nM). Medium, target cells, and constructs were replenished on day 4, followed by flow cytometric analysis on day 5 (**F**) or day 7 (**G**). (**F**) Representative T cell proliferation profiles (left panel) and division index of proliferating T cells (n=4 donors) calculated using FlowJo software (right panel). (**G**) Representative tSNE plots and central memory T cell counts were assessed on day 7 (n=7 donors).

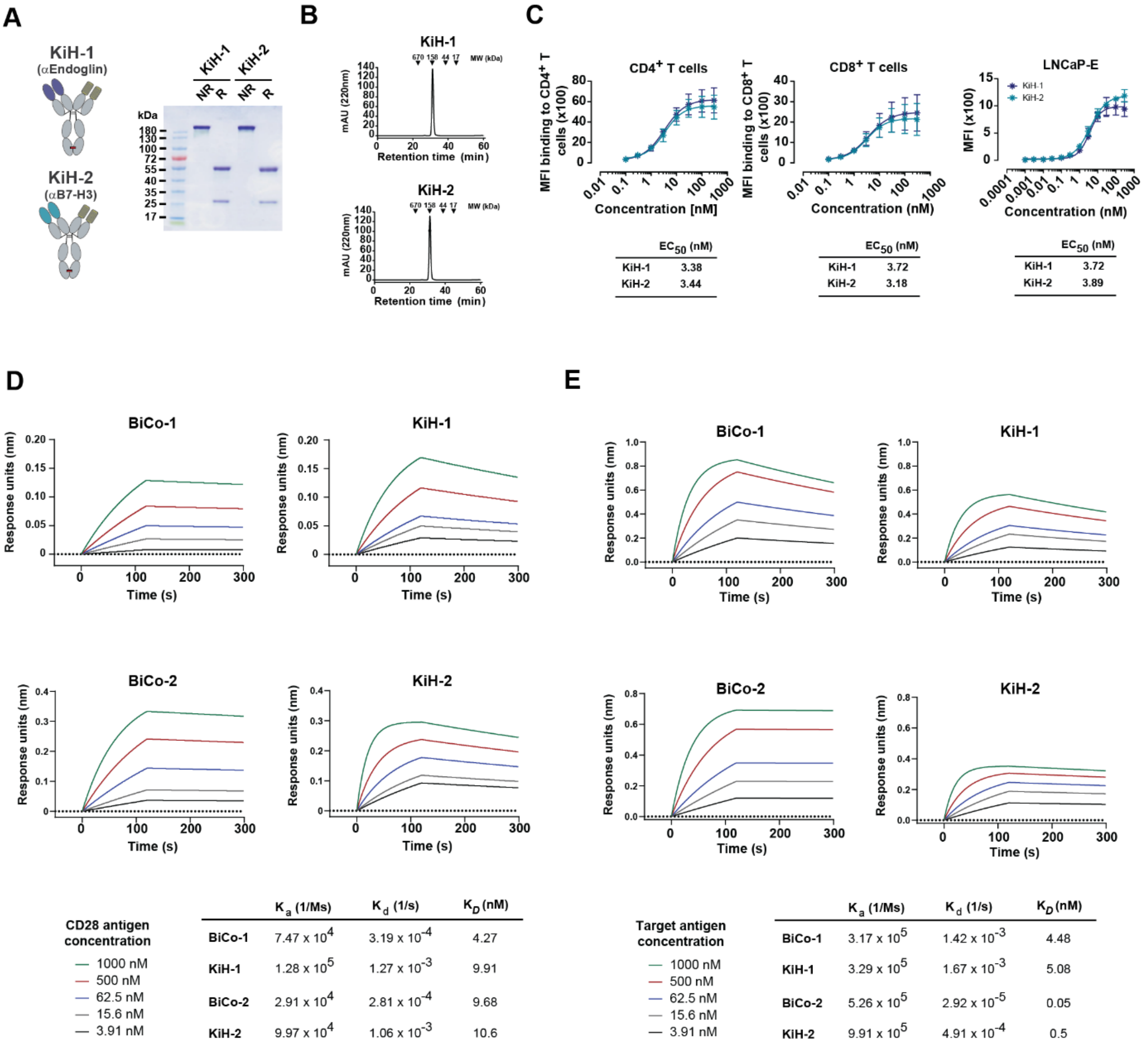


**Supplemental Figure 6 | Generation and characterization of knob-into-hole (KiH) bispecific CD28 antibodies and comparison of binding characteristics with BiCos**

(**A**) Schematic representation of knob-into-hole bispecific costimulatory antibodies (KiH-1, endoglin×CD28; KiH-2, B7-H3×CD28) and SDS-PAGE analysis under non-reducing (NR) and reducing (R) conditions, confirming correct molecular assembly.

(**B**) Analytical size-exclusion chromatography of KiH-1 and KiH-2 performed on a Superdex S200 Increase 10/300 GL column.

(**C**) Binding of the indicated KiH constructs to CD28-expressing $CD4^+$ and $CD8^+$ T-cells (n=4 donors) and to LNCaP-E tumor cells as assessed by flow cytometry. Data are shown as mean ± SD. $EC_{50}$ values were calculated using GraphPad Prism software.

(**D-E**) Binding kinetics of BiCo molecules and the corresponding KiH constructs to recombinant human CD28 (**D**) and the target antigens endoglin and B7-H3 (**E**) determined by Biolayer interferometry (Octet). Kinetic parameters were derived from global fitting, and apparent affinities were calculated.

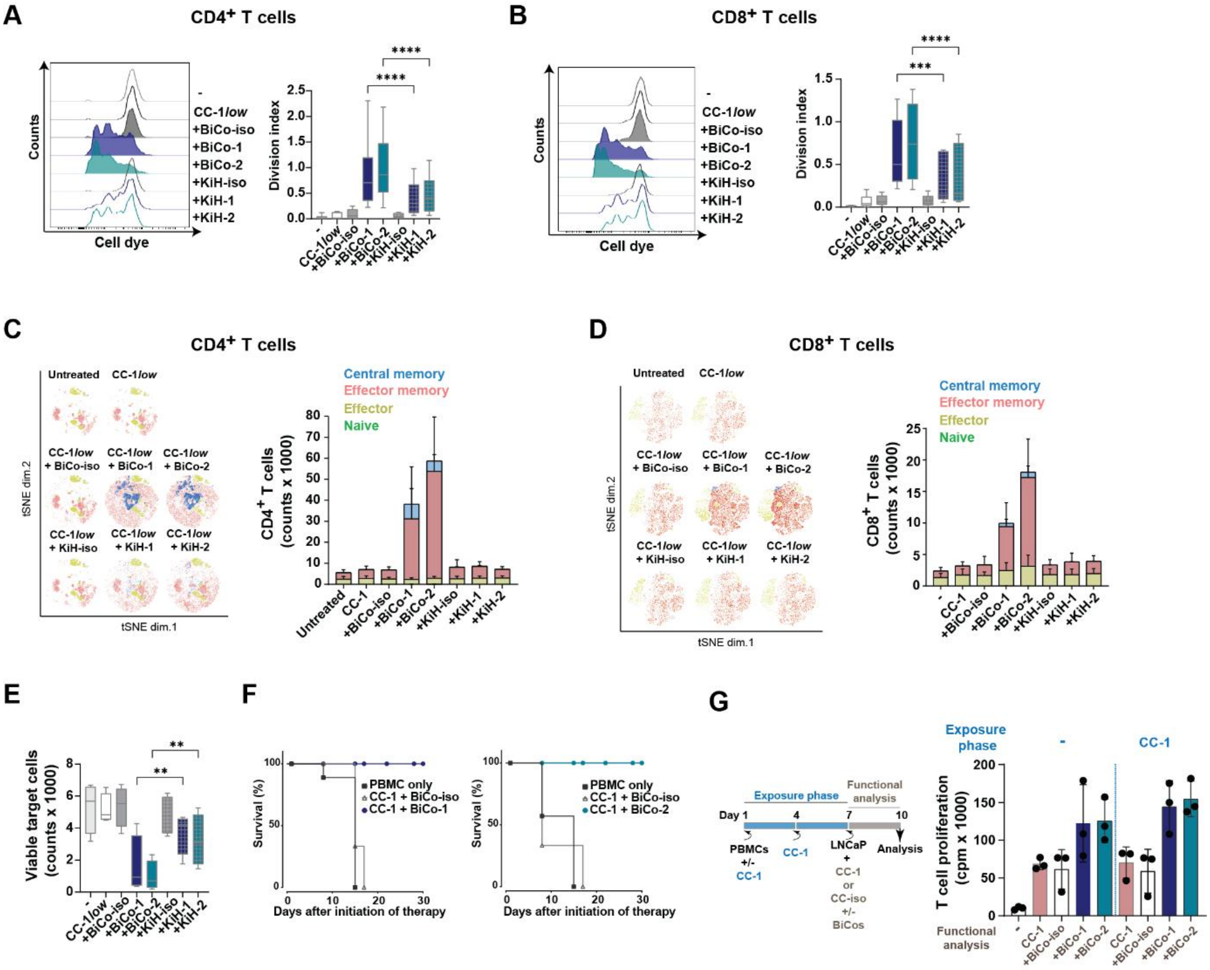

**Supplemental Figure 7 | BiCo-1 and BiCo-2 promote CC-1–dependent T-cell proliferation, differentiation, and tumor-cell killing.**

**(A-E)** CellTrace Violet–labeled monocyte-depleted PBMC were cultured with LNCaP-E cells at an E:T ratio of 1:1 with CC-1 (25 pM, CC-1low) and BiCo/KiH (both at 0.5 nM) as indicated. Medium, target cells, and constructs were replenished on day 4, followed by flow cytometric analysis. (**A, B**) Representative proliferation profiles (left panels) on day 5. Division index (n=6 donors) of proliferating CD4$^{+}$ T cells and CD8$^{+}$ T cells was calculated using FlowJo (right panels). (**C, D**) Representative tSNE plots (left) and T cell subset counts (right) for CD4$^{+}$ (**C**) and CD8$^{+}$ T cells (**D**) were determined on day 7 (n=6 donors). (**E**) Tumor cell killing as assessed at day 7 (n=4 donors).

(**F**) Kaplan-Meier survival curve following therapy initiation using animal sacrifice as the terminal event. The results relate to the in vivo experiment depicted in the Figure 2O.

(**G**) PBMCs (n = 3) were cultured in the presence or absence of CC-1 (1 nM); medium with or without CC-1 was replenished on day 4. On day 7, PBMCs were washed and incubated with LNCaP-E cells at an E:T ratio of 1:1 in the presence of CC-1 (1 nM) alone or in combination with the indicated BiCo constructs (0.5 nM). On day 10, T cell proliferation was assessed by $^{3}$H-thymidine incorporation.

**Supplemental table 1: TotalSeq-C Hashtag Oligonucleotide-Conjugated Antibodies**

| Antibody | Barcode Sequence | Catalog No. | Manufacturer |
|---|---|---|---|
| TotalSeq™-C0251 anti-human Hashtag 1 Antibody | GTCAACTCTTTAGCG | 394661 | BioLegend |
| TotalSeq™-C0252 anti-human Hashtag 2 Antibody | TGATGGCCTATTGGG | 394663 | BioLegend |
| TotalSeq™-C0253 anti-human Hashtag 3 Antibody | TTCCGCCTCTCTTTG | 394665 | BioLegend |
| TotalSeq™-C0254 anti-human Hashtag 4 Antibody | AGTAAGTTCAGCGTA | 394667 | BioLegend |
| TotalSeq™-C0255 anti-human Hashtag 5 Antibody | AAGTATCGTTTCGCA | 394669 | BioLegend |

**Supplemental table 2: Directly Conjugated Antibodies Used for Flow Cytometry**

| **Antibody** | **Fluorochrome** | **Clone** | **Catalog No.** | **Manufacturer** |
|---|---|---|---|---|
| Human CD4 | APC/Cy7 | RPA-T4 | 300518 | BioLegend |
| Human CD4 | Pacific Blue | OKT4 | 317429 | BioLegend |
| Human CD8a | FITC | RPA-T8 | 301006 | BioLegend |
| Human CD25 | PE | BC96 | 302606 | BioLegend |
| Human CD27 | AF700 | MT271 | 356416 | BioLegend |
| Human CD28 | Pe-Fire810 | CD28.2 | 302971 | BioLegend |
| Human CD45RO | PE/Cy7 | UCHL1 | 304230 | BioLegend |
| Human CD57 | Pe-Dazzle | HNK1 | 359620 | BioLegend |
| Human CD62L | Pacific Blue | DREG-56 | 304826 | BioLegend |
| Human CD62L | BV421 | DREG-56 | 304828 | BioLegend |
| Human CD69 | APC/Cy7 | FN50 | 310914 | BioLegend |
| Human EpCAM | PE/Cy7 | 9C4 | 324222 | BioLegend |
| Human TIGIT | KiraviaBlue520 | A15153G | 372732 | BioLegend |
| Human PD-1 | BV480 | EH12.1 | 566112 | BD |
| Human KLRG1 | PerCP/Fire | SA231A2 | 367746 | BioLegend |
| Human LAG3 | BUV805 | 3DS223H | 368-2239-42 | Thermo Fisher |
| Human TIM3 | BV605 | F38-2E2 | 345018 | BioLegend |
